## Supplementary material for "A tale of two birds: cognitive simplicity drives collective route improvements in homing pigeons": SI Appendix

**This PDF file includes:**

Supporting text

Figures S1

Tables S1-S6

### SUPPLEMENTARY INFORMATION

#### Model Evaluation

1. **Route Efficiency Model:** We used `Gen` (generation), `strategy` (0 for experimental and 1-7 for simulated data) and `bird_ID` as our predictor variables with route efficiency as our response variable. Broadly we tested fitting different model formulations including random effects and variable relationship. The code for the models we formulated are available in the `pigeon_replace_model_performance.R` file. We used AIC to select the best fitting model:

| **Model** | **df** | **AIC** |
|---|---|---|
| Linear Model | 41 | -2141657 |
| Linear Mixed Effect Model (random effect bird ID) | 42 | -2280167 |
| Linear Mixed Effect Model (random effect bird ID & Generation) | 42 | -2281106 |
| Beta Regression Model | 41 | -2388197 |
| Beta Regression Model (random effect bird ID) | 42 | -2526965 |
| Beta Regression Model (random effect bird ID & Generation) | 42 | -2527103 |

The lowest AIC value was achieved by our beta regression model (See *Methods* in the main text for details). The model diagnostics are:

| **Metric** | **Value** |
|---|---|
| Fixed Effect Variance | 0.0425 |
| Random Variance | 0.1080 |
| Residual Variance | 0.00048 |
| Marginal $R^2$ | 0.2815 |
| Conditional $R^2$ | 0.9968 |
| Intraclass Correlation Coefficient (ICC) | 0.7154 |

2. **Social Weight Model:** We used similar predictor variables as the prior model and tested different model formulations, with social weight of the experienced bird as our response variable. The AIC results were:

| **Model** | **df** | **AIC** |
|---|---|---|
| Linear Model | 21 | -18914.90 |
| Linear Mixed Effect Model (random effect bird ID) | 22 | -81533.43 |
| Linear Mixed Effect Model (random effect bird ID & Generation) | 22 | -81967.68 |

The best fitting model was a mixed effect model with our random effect terms. Model diagnostics:

| Metric | Value |
|---|---|
| Fixed Effect Variance | 0.0038 |
| Random Variance | 0.0257 |
| Residual Variance | 0.0429 |
| Marginal $R^2$ | 0.0527 |
| Conditional $R^2$ | 0.4075 |
| Intraclass Correlation Coefficient (ICC) | 0.3548 |

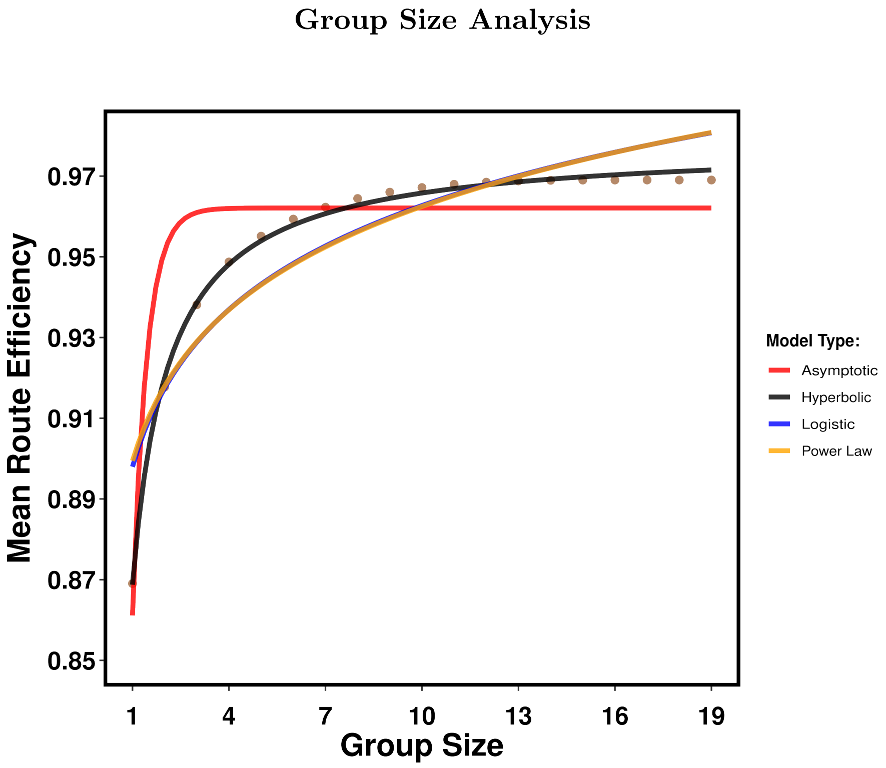


Figure S1. The figure illustrates various functional models fitted to the mean route efficiency values for a given group size. A post-hoc model comparison was conducted using AIC to determine the best-fitting model: hyperbolic (df = 3, AIC = -189.8), logistic (df = 3, AIC = -114.6), power law (df = 3, AIC = -113.5), and asymptotic (df = 3, AIC = -110.5). Among these, the hyperbolic model provided the best fit. Note that the power law and logistic curves in the figure are nearly identical.

| **Contrast** | **Odds Ratio** | **SE** | **CI (95%)** | **p-value** | |
|---|---|---|---|---|---|
| **Averaging** | 1.05 | 0.07 | [0.88, 1.25] | 0.93 | |
| **Experienced Bird Weighting** | 1.13 | 0.08 | [0.94, 1.34] | 0.34 | |
| **Maximize Generation** | 1.15 | 0.08 | [0.96, 1.37] | 0.19 | |
| **Better Bird Weighting** | **1.37** | **0.09** | **[1.15, 1.63]** | **<.001** | *** |
| **All-or-Nothing** | 0.99 | 0.07 | [0.84, 1.19] | 1.00 | |
| **Maximize Flight** | **1.55** | **0.11** | **[1.30, 1.86]** | **<.001** | *** |
| **Maximize Time Steps** | **1.26** | **0.08** | **[1.06, 1.51]** | **<.01** | ** |

Table S1. The table represents contrasts between the estimated route efficiencies of each strategy and the experimental data holding generation constant (at its mean value). The contrasts were conducted on the log odds ratio scale relative to the experimental data. Confidence intervals (CIs) for contrasts were calculated at 95% interval. To account for multiple comparisons with the experimental data (seven tests per generation), Dunnett's correction was applied to both CIs and p-values. Significance levels are denoted as follows: * $p < 0.05$, ** $p < 0.01$, *** $p < 0.001$.

| **Contrast** | **Mean Difference** | **SE** | **CI (95%)** | **p-value** | |
|---|---|---|---|---|---|
| **Averaging** | 0.003 | 0.03 | [-0.08, 0.09] | 0.99 | |
| **Experienced Bird Weighting** | **0.14** | **0.03** | **[0.05, 0.23]** | **<.001** | *** |
| **Maximize Generation** | **0.18** | **0.04** | **[0.09, 0.27]** | **<.001** | *** |
| **Better Bird Weighting** | **0.17** | **0.04** | **[0.06, 0.29]** | **<.001** | *** |
| **All-or-Nothing** | **0.23** | **0.04** | **[0.11, 0.34]** | **<.001** | *** |
| **Maximize Flight** | **0.23** | **0.04** | **[0.11, 0.34]** | **<.001** | *** |
| **Maximize Time Steps** | **0.12** | **0.04** | **[0.00, 0.23]** | **0.04** | * |

Table S2. The table represents contrasts between the estimated social weights of the experienced bird for each strategy and the experimental data holding generation constant (at its mean value). The contrasts were conducted on the response scale relative to the experimental data. CIs for contrasts were calculated at the 95% interval. To account for multiple comparisons with the experimental data (seven tests per generation), Dunnett's correction was applied to both CIs and p-values. Significance levels are denoted as follows: * $p < 0.05$, ** $p < 0.01$, *** $p < 0.001$.

| Contrast | Generation | Odds Ratio | SE | CI (95%) | p-value | |
|---|---|---|---|---|---|---|
| | 1 | 1.12 | 0.17 | [0.75, 1.67] | 0.91 | |
| **Averaging** | 2 | 0.98 | 0.14 | [0.67, 1.43] | 1.00 | |
| | 3 | 1.29 | 0.18 | [0.90, 1.85] | 0.30 | |
| | 4 | 0.90 | 0.14 | [0.60, 1.35] | 0.94 | |
| | 5 | 1.05 | 0.16 | [0.70, 1.58] | 0.99 | |
| | 1 | 1.12 | 0.17 | [0.75, 1.67] | 0.91 | |
| **Experienced Bird Weighting** | 2 | 0.98 | 0.14 | [0.67, 1.43] | 1.00 | |
| | 3 | 1.37 | 0.19 | [0.95, 1.96] | 0.13 | |
| | 4 | 1.02 | 0.16 | [0.68, 1.53] | 1.00 | |
| | 5 | 1.23 | 0.19 | [0.82, 1.86] | 0.60 | |
| | 1 | 1.12 | 0.17 | [0.75, 1.67] | 0.91 | |
| **Maximize Generation** | 2 | 0.98 | 0.14 | [0.67, 1.43] | 1.00 | |
| | 3 | 1.40 | 0.19 | [0.97, 2.03] | 0.08 | |
| | 4 | 1.08 | 0.17 | [0.71, 1.63] | 0.74 | |
| | 5 | 1.36 | 0.21 | [0.89, 2.07] | 0.22 | |
| | 1 | 1.12 | 0.17 | [0.75, 1.67] | 0.91 | |
| **Better Bird Weighting** | 2 | 1.15 | 0.16 | [0.78, 1.69] | 0.83 | |
| | **3** | **1.66** | **0.23** | **[1.16, 2.39]** | **<.01** | ** |
| | 4 | 1.27 | 0.20 | [0.84, 1.90] | 0.48 | |
| | **5** | **1.57** | **0.24** | **[1.05, 2.37]** | **0.02** | * |
| | 1 | 1.12 | 0.17 | [0.75, 1.67] | 0.91 | |
| **All-or-Nothing** | 2 | 0.94 | 0.13 | [0.64, 1.38] | 1.00 | |
| | 3 | 1.23 | 0.17 | [0.85, 1.78] | 0.14 | |
| | 4 | 0.88 | 0.14 | [0.58, 1.34] | 0.89 | |
| | 5 | 1.06 | 0.16 | [0.70, 1.61] | 0.99 | |
| | 1 | 1.12 | 0.17 | [0.75, 1.67] | 0.91 | |
| **Maximize Flight** | 2 | 1.23 | 0.18 | [0.83, 1.79] | 0.54 | |
| | **3** | **1.89** | **0.26** | **[1.32, 2.72]** | **<.001** | *** |
| | **4** | **1.51** | **0.23** | **[1.00, 2.27]** | **0.046** | * |
| | **5** | **1.96** | **0.30** | **[1.30, 2.94]** | **<.001** | *** |
| | 1 | 1.12 | 0.17 | [0.75, 1.67] | 0.91 | |
| **Maximize Time Steps** | 2 | 1.07 | 0.15 | [0.73, 1.57] | 0.76 | |
| | **3** | **1.54** | **0.21** | **[1.07, 2.22]** | **0.01** | * |
| | 4 | 1.18 | 0.18 | [0.79, 1.78] | 0.76 | |
| | 5 | 1.49 | 0.23 | [0.99, 2.24] | 0.06 | |

Table S3. The table represents contrasts derived from the Estimated Marginal Means (EMMs) of the beta regression model applied to the route efficiency data as our response variable. The contrasts were conducted on the log odds ratio scale relative to the experimental data. CIs for contrasts were calculated at 95% interval. To account for multiple comparisons with the experimental data (seven tests per generation), Dunnett's correction was applied to both CIs and p-values. Significance levels are denoted as follows: * $p < 0.05$, ** $p < 0.01$, *** $p < 0.001$.

| Contrast | Generation | Mean Difference | SE | CI (95%) | p-value |
|---|---|---|---|---|---|
| | 2 | -0.03 | 0.06 | [-0.19, 0.14] | 0.98 |
| **Averaging** | 3 | 0.05 | 0.06 | [-0.12, 0.22] | 0.92 |
| | 4 | -0.05 | 0.07 | [-0.24, 0.14] | 0.95 |
| | 5 | 0.04 | 0.08 | [-0.17, 0.24] | 0.98 |
| | 2 | 0.11 | 0.06 | [-0.05, 0.28] | 0.38 |
| **Experienced Bird Weighting** | **3** | **0.19** | **0.06** | **[0.02, 0.38]** | **0.02** * |
| | 4 | 0.09 | 0.07 | [-0.09, 0.28] | 0.64 |
| | 5 | 0.17 | 0.08 | [-0.03, 0.38] | 0.13 |
| | 2 | -0.03 | 0.06 | [-0.20, 0.14] | 0.98 |
| **Maximize Generation** | **3** | **0.22** | **0.06** | **[0.04, 0.39]** | **<.01** ** |
| | **4** | **0.20** | **0.08** | **[0.01, 0.39]** | **0.03** * |
| | **5** | **0.33** | **0.08** | **[0.13, 0.54]** | **<.001** *** |
| | 2 | -0.03 | 0.08 | [-0.23, 0.17] | 0.99 |
| **Better Bird Weighting** | **3** | **0.22** | **0.08** | **[0.00, 0.44]** | **0.05** * |
| | 4 | 0.19 | 0.09 | [-0.04, 0.43] | 0.17 |
| | **5** | **0.29** | **0.09** | **[0.05, 0.54]** | **0.01** * |
| | 2 | -0.03 | 0.08 | [-0.26, 0.20] | 0.99 |
| **All-or-Nothing** | **3** | **0.26** | **0.08** | **[0.03, 0.48]** | **0.01** * |
| | **4** | **0.26** | **0.09** | **[0.02, 0.49]** | **0.02** * |
| | **5** | **0.39** | **0.09** | **[0.14, 0.64]** | **<.001** *** |
| | 2 | -0.03 | 0.09 | [-0.26, 0.20] | 0.99 |
| **Maximize Flight** | **3** | **0.27** | **0.08** | **[0.04, 0.49]** | **0.01** * |
| | **4** | **0.26** | **0.09** | **[0.05, 0.49]** | **0.03** * |
| | **5** | **0.39** | **0.09** | **[0.14, 0.63]** | **<.001** *** |
| | 2 | -0.03 | 0.09 | [-0.26, 0.20] | 0.99 |
| **Maximize Time Steps** | 3 | 0.17 | 0.08 | [-0.05, 0.39] | 0.23 |
| | 4 | 0.14 | 0.09 | [-0.10, 0.37] | 0.48 |
| | **5** | **0.26** | **0.09** | **[0.00, 0.50]** | **0.04** * |

Table S4. The table represents contrasts derived from the Estimated Marginal Means (EMMs) of the linear mixed model applied to the social weights of the experienced bird as our response variable. The contrasts were conducted on the response scale with mean difference relative to the experimental data. CIs for contrasts were calculated at 95% interval. To account for multiple comparisons with the experimental data (seven tests per generation), Dunnett's correction was applied to both CIs and p-values. Significance levels are denoted as follows: * $p < 0.05$, ** $p < 0.01$, *** $p < 0.001$.

| Contrast | Generation | Odds Ratio | CI (95%) | p-value | |
|---|---|---|---|---|---|
| | 1 | 1.09 | [0.51, 2.35] | 1.0 | |
| **Averaging** | 2 | 1.01 | [0.49, 2.07] | 1.0 | |
| | 3 | 1.47 | [0.72, 3.00] | 0.53 | |
| | 4 | 0.86 | [0.45, 1.65] | 0.95 | |
| | 5 | 1.01 | [0.48, 2.17] | 1.0 | |
| | 1 | 1.08 | [0.51, 2.33] | 1.0 | |
| **Experienced Bird Weighting** | 2 | 0.97 | [0.46, 2.04] | 1.0 | |
| | 3 | 1.54 | [0.76, 3.09] | 0.42 | |
| | 4 | 0.97 | [0.54, 1.76] | 1.0 | |
| | 5 | 1.21 | [0.61, 2.39] | 0.92 | |
| | 1 | 1.09 | [0.51, 2.34] | 1.0 | |
| **Maximize Generation** | 2 | 1.01 | [0.49, 2.06] | 1.0 | |
| | 3 | 1.58 | [0.79, 3.14] | 0.35 | |
| | 4 | 1.02 | [0.58, 1.79] | 1.0 | |
| | 5 | 1.32 | [0.70, 2.50] | 0.71 | |
| | 1 | 1.09 | [0.51, 2.34] | 1.0 | |
| **Better Bird Weighting** | 2 | 1.20 | [0.60, 2.43] | 0.93 | |
| | **3** | **1.96** | **[1.08, 3.55]** | **0.01** | * |
| | 4 | 1.26 | [0.83, 1.91] | 0.52 | |
| | 5 | 1.61 | [0.96, 2.69] | 0.08 | |
| | 1 | 1.09 | [0.51, 2.35] | 1.0 | |
| **All-or-Nothing** | 2 | 0.99 | [0.52, 1.89] | 1.0 | |
| | 3 | 1.44 | [0.82, 2.51] | 0.38 | |
| | 4 | 0.87 | [0.61, 1.22] | 0.74 | |
| | 5 | 1.07 | [0.69, 1.65] | 0.99 | |
| | 1 | 1.09 | [0.51, 2.34] | 1.0 | |
| **Maximize Flight** | 2 | 1.12 | [0.62, 2.65] | 0.85 | |
| | **3** | **2.21** | **[1.19, 4.12]** | **<.01** | ** |
| | **4** | **1.51** | **[1.00, 2.27]** | **0.047** | * |
| | **5** | **2.01** | **[1.26, 3.23]** | **<.001** | *** |
| | 1 | 1.09 | [0.51, 2.34] | 1.0 | |
| **Maximize Time Steps** | 2 | 1.16 | [0.55, 2.27] | 0.99 | |
| | 3 | 1.80 | [0.96, 3.36] | 0.08 | |
| | 4 | 1.16 | [0.74, 1.80] | 0.87 | |
| | 5 | 1.49 | [0.89, 2.50] | 0.20 | |

Table S5. Summary of bootstrap contrasts for different strategies relative to the experimental control for the route efficiency data. The table reports odds ratios, 95% CIs and adjusted p-values. Multiple comparison adjustments were made using the Dunnett's method. Significance levels: * $p < 0.05$, ** $p < 0.01$, *** $p < 0.001$.

| Contrast | Generation | Mean Difference | CI (95%) | p-value | |
|---|---|---|---|---|---|
| **Averaging** | 2 | -0.03 | [-0.24, 0.18] | 0.99 | |
| | 3 | 0.05 | [-0.14, 0.23] | 0.93 | |
| | 4 | -0.05 | [-0.34, 0.24] | 0.98 | |
| | 5 | 0.04 | [-0.24, 0.31] | 0.99 | |
| **Experienced Bird Weighting** | 2 | 0.11 | [-0.10, 0.32] | 0.58 | |
| | **3** | **0.19** | **[0.01, 0.37]** | **0.041** | * |
| | 4 | 0.09 | [-0.20, 0.38] | 0.88 | |
| | 5 | 0.18 | [-0.10, 0.45] | 0.37 | |
| **Maximize Generation** | 2 | -0.03 | [-0.24, 0.18] | 0.99 | |
| | **3** | **0.22** | **[0.03, 0.49]** | **0.01** | * |
| | 4 | 0.20 | [-0.08, 0.49] | 0.29 | |
| | **5** | **0.34** | **[0.06, 0.61]** | **<.01** | ** |
| **Better Bird Weighting** | 2 | -0.03 | [-0.36, 0.31] | 1.00 | |
| | 3 | 0.22 | [0.00, 0.49] | 0.16 | |
| | 4 | 0.19 | [-0.13, 0.52] | 0.47 | |
| | **5** | **0.30** | **[0.06, 0.61]** | **<.01** | ** |
| **All-or-Nothing** | 2 | -0.03 | [-0.52, 0.47] | 1.00 | |
| | 3 | 0.26 | [-0.17, 0.68] | 0.44 | |
| | 4 | 0.26 | [-0.21, 0.73] | 0.53 | |
| | 5 | 0.40 | [-0.04, 0.84] | 0.09 | |
| **Maximize Flight** | 2 | -0.03 | [-0.32, 0.26] | 1.00 | |
| | **3** | **0.27** | **[0.01, 0.53]** | **0.04** | * |
| | 4 | 0.26 | [-0.07, 0.59] | 0.21 | |
| | **5** | **0.39** | **[0.08, 0.70]** | **<.01** | ** |
| **Maximize Time Steps** | 2 | -0.03 | [-0.35, 0.29] | 1.00 | |
| | 3 | 0.16 | [-0.12, 0.45] | 0.46 | |
| | 4 | 0.14 | [-0.23, 0.50] | 0.82 | |
| | 5 | 0.26 | [-0.10, 0.61] | 0.28 | |

Table S6. Summary of bootstrap mean difference contrasts for different strategies relative to the experimental control for the social weights data. The table reports mean differences, 95% CIs (adjusted for multiple comparisons), and adjusted p-values using the Dunnett's method. Significance levels: * $p < 0.05$, ** $p < 0.01$, *** $p < 0.001$.
